# Pathways of human development threaten biomes’ protection and their remaining natural vegetation

**DOI:** 10.1101/776443

**Authors:** Isabel M.D. Rosa, Carlos A. Guerra

## Abstract

Protected areas have been one of the most commonly applied conservation tools to prevent ecosystem degradation. International conservation targets have been created to incentivize widespread expansion of protected area networks, but this call might clash with expected future land use change. Here we investigated how future land use trajectories (2015-2090), representing a wide range of plausible future scenarios would impact the remaining areas of primary vegetation under different protection levels across the world’s biomes. We then highlight areas under greater risk of conflict between conservation (highly protected) and land use expansion (high projected change), and areas where these two can better co-exist (lower protection with high projected change and/or high protection with low projected change).

While the most positive pathway of development led to the least loss of primary vegetation globally, this was not observed in all biomes. Further, we found no significant correlation between existing extent of protection and average proportion of vegetation loss. Mediterranean Forests, Woodlands & Scrub had the largest projected loss occurring in the highest protected areas. Tropical Forests in Central Africa and the Boreal Forests of North Euro-Asia and Canada emerge as the areas where most projected change occurs, and existing protection is still low. Areas in India and Southeast Asia emerge as potential areas for intervention as they have significant projected loss of primary vegetation, and considerably low protection.

Our results can help inform policy and decision-makers to prevent such conflicts and support the development of management actions. These policy and management actions should target conservation in areas under expected great pressure of change with high ecological value (e.g., composed mainly by primary vegetation), but still not protected. This study also opens the discussion to the future of current protected areas and to the potential to expand the existing network of protected areas.

## Introduction

Humans have been degrading and shaping landscapes worldwide for many centuries (Ellis & Ramankutty, 2008). In fact, in 1700, nearly half of the terrestrial biosphere was wild, whereas by 2000, the majority of the terrestrial ecosystems was already converted into agricultural lands and settlements, leaving less than 20% of semi-natural areas and only a quarter left wild (Ellis, Klein Goldewijk, Siebert, Lightman, & Ramankutty, 2010). This trend of human modification of landscapes is expected to continue as human population keeps increasing and, as a consequence, so does the demand for agricultural and forest products (Boserup, 2017). Moreover, as humans convert natural habitats (Gibbs et al., 2010), the world’s biomes and ecoregions become more degraded, jeopardizing these as habitats for species and hampering the benefits people derive from them (Díaz et al., 2018; Hoekstra, Boucher, Ricketts, & Roberts, 2005). A central challenge of achieving sustainability is, therefore, how to preserve natural ecosystems while enhancing food production (Lambin & Meyfroidt, 2011).

Protected areas have long been used as important conservation tools to prevent ecosystem degradation and preserve biodiversity and ecosystem services vital to sustain human livelihoods (Watson, Dudley, Segan, & Hockings, 2014). As such, the number of protected areas has increased greatly since the 1990s (Anthamatten & Hazen, 2015). However, the overall coverage of these areas is still rather low, i.e. roughly 12-13% (Anthamatten & Hazen, 2015; Brooks, Da Fonseca, & Rodrigues, 2004; Jenkins & Joppa, 2009), reducing to 9.3% when considering well-connected protected areas (Saura, Bastin, Battistella, Mandrici, & Dubois, 2017). There is, nonetheless, international pressure to increase this coverage, especially by the establishment of international conservation targets, such as the Aichi Targets, specifically Target 11, which states that by 2020 at least 17% of terrestrial areas are conserved through well-connected systems of protected areas (https://www.cbd.int/sp/targets/). This call to expand protected areas might clash with the expected expansion of agricultural lands for food production and other types of land use change.

The relationship between the effectiveness and the placement of these protected areas has been a great source of debate. Claims have been made that protected areas are often located in remote areas (Joppa & Pfaff, 2009), isolated and with low population densities (Baldi, Texeira, Martin, Grau, & Jobbágy, 2017), thus using the landscape characteristics (higher slope, further from roads and cities) to explain why they suffer less degradation (Schulze et al., 2018). Others, however, have shown the ‘pulling’ effect of these areas, with land cover change occurring closer to protected areas than in more distant unprotected lands (Guerra et al. in review). Simultaneously, it has been shown that pressure on protected areas has increased over time (Geldmann, Joppa, & Burgess, 2014), particularly in developing countries threatened by resource (over)exploitation (Schulze et al., 2018). Nonetheless, there is mounting evidence that protected areas have a positive influence in maintaining the natural habitats (Paiva, Brites, & Machado, 2015), and on their ability to sustain higher levels of biodiversity (Gray et al., 2016; Thomas & Gillingham, 2015); with the differences mostly attributable to differences in land use between protected and unprotected sites (Gray et al., 2016).

Thus, to maximize conservation outcomes, it is crucial to identify areas with the greatest potential to expand protected areas. Nevertheless, this comes with the risk of ineffective outcomes due to land use change and uncoordinated actions between countries (Pouzols et al., 2014). Previous studies have shown that under different scenarios of land use change it might become infeasible to achieve the 17% of terrestrial land protected, which when combined with increasing land use change threatens a high number of species (Pouzols et al., 2014). Also, a continued decline of primary vegetation lands within the areas surrounding protected areas is expected thus leading to an increasingly heterogeneous matrix of primary and human-modified landscapes (Beaumont & Duursma, 2012).

For the foreseeable future, the fate of terrestrial ecosystems and the species they support will continue to be intertwined with human systems, as most of the remaining natural areas are now embedded within anthropogenic mosaics of land use. However, the rate and location of land use change required to meet the demand for commodities are highly uncertain as it depends on the trajectories of development that might unfold in the future. In this regard, a set of Shared Socio-economic Pathways (SSPs), associated with the Representative Concentration Pathways (RCPs), have been developed by the climate science community (O’Neill et al., 2017, 2014; Van Vuuren et al., 2011). Working under the Intergovernmental Panel on Climate Change (IPCC) auspices, these SSPs and RCPs describe different scenarios of human development trajectories that would result in different climate futures based on land use change projections and greenhouse gas emissions over the 21^st^ century (Popp et al., 2017; Riahi et al., 2017). In particular, the SSPs explore a wide range of scenarios on climate change mitigation and adaptation, on technological improvements, on economic developments and population growth, covering a range of futures from a sustainable and environmentally-friendly world (SSP1) to a world continued to be dominated by fossil fuels (SSP5) (Riahi et al., 2017). Each SSP has its own storyline with associated projected land use change (Table 1), as described in (Popp et al., 2017).

**Table 1.**
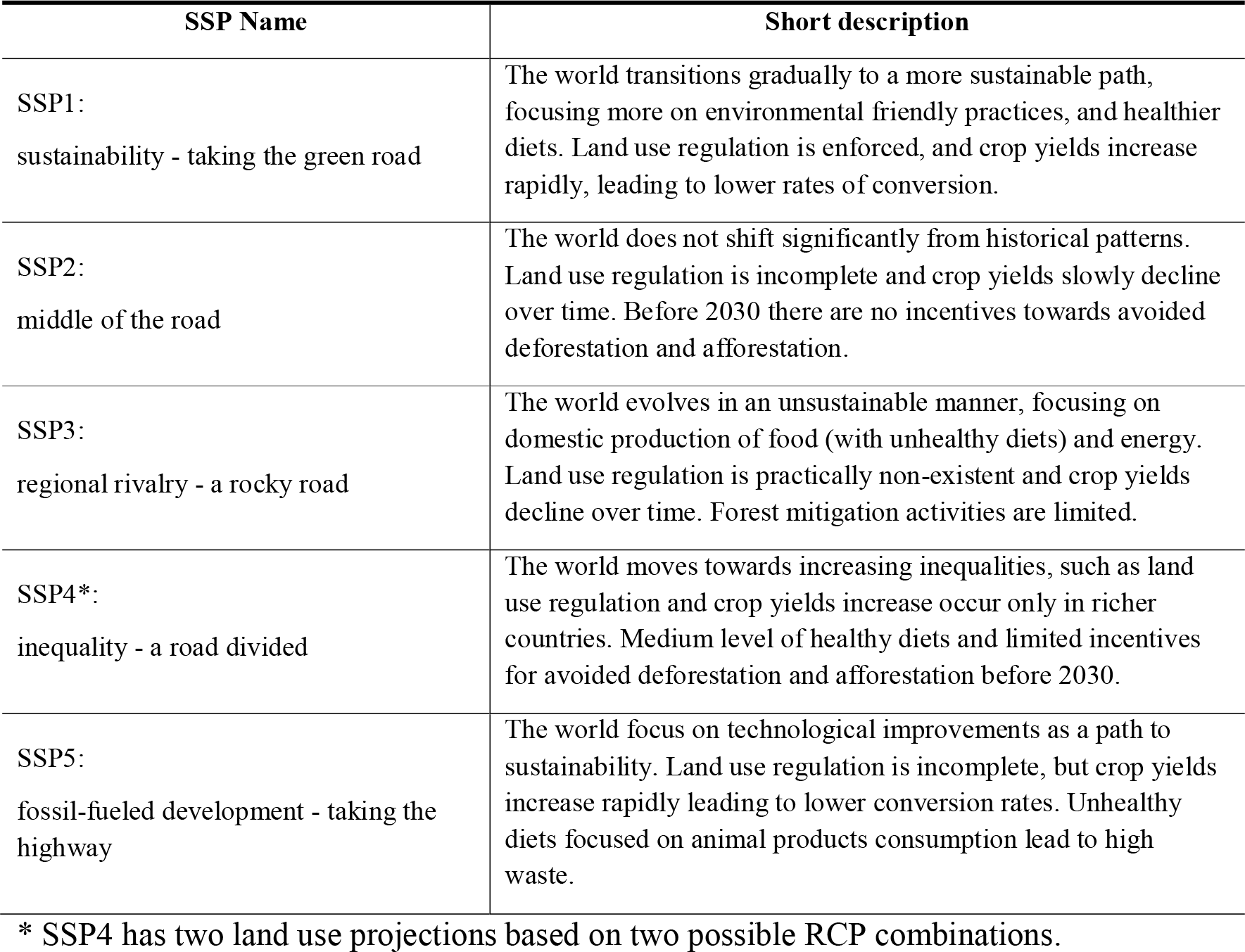
Short description of the five Shared Socio-economic Pathways (SSPs) storylines with particular focus on the associated consequences for land use change (adapted from Popp et al. 2017). For a detailed description of the narratives of each SSP, please see Popp et al. 2017 and Riahi et al. 2017.

As a major driver of biodiversity and ecosystem services change, with significant impacts on climate and ultimately human well-being, is thus important to understand how current conservation areas might be impacted by these projections of future land use change. Therefore, the main objective of this study was to investigate how future land use trajectories, representing a wide range of plausible future scenarios (the five SSPs), would impact areas of primary vegetation under different protection, across the world’s biomes from 2015 through 2090. With such analysis, we aimed to highlight areas under greater risk of conflict between conservation (highly protected) and land use expansion (high projected change), and areas where these two can better co-exist (lower protection with high projected change and/or high protection with low projected change). Such results could help inform policy and decision-makers to prevent such conflicts and support the development of management actions targeting conservation in areas under expected great pressure of change and high ecological value (e.g., composed mainly by primary vegetation), but still not protected (i.e., potential areas to expand existing network of protected areas).

## Methods

### Input Data and Sources

We used the land use projections provided by the dataset of the Land Use Harmonized v2.0 project (http://luh.umd.edu/) (Hurtt et al., 2011; Hurtt et al., 2016). The dataset, which was produced within the context of the World Climate Research Program Coupled Model Intercomparison Project (CMIP6), contains a harmonized set of land use scenarios that are consistent between historical reconstructions and future projections. In detail, it contains annual land use maps, produced by different integrated assessment models (IAMs) for each SSP, from 2015 through 2100 at 0.25° resolution, with the proportion of each pixel covered by each one of 12 land use classes (Table S1). In this study, we focused specifically on the loss of primary vegetation land (both forested and non-forested) given that protected areas are mainly implemented to protect pristine environments and not human-modified lands (Baldi et al., 2017; Paiva et al., 2015). The resolution of the land use time series dataset determined the spatial unit of analysis, and for each SSP we obtained a different time series of projected land use change according to the assumptions of each pathway (Table 1, see details in Riahi et al., 2017), and the model used to spatialize these assumptions (Popp et al., 2017). As we intended to focus our analysis only on the loss of primary vegetation, we aggregated the original land use classes into two: primary and modified as detailed in Table S1.

One limitation of our study is the fact that the categories of land use provided by the LUH2 project are spatially and descriptively coarse. Although these categories have greatly improved since LUH1 (Beaumont & Duursma, 2012), these still do not allow us to discriminate exactly the land use matrix within each 0.25 × 0.25° grid cell. This means that our analysis is blind to the detailed spatial configuration of loss in primary vegetation, i.e., whether a projected 10% loss in primary vegetation is adjacent to existing loss, or spread homogeneously across the grid cell.

Furthermore, we used the entire geodatabase of the World Database of Protected Areas (Brooks et al., 2004; Dubois et al., 2016), as of October 2018, to obtain the geographic location of all current protected areas in the world. From this dataset we produced a raster with the same extent and cell size as the land use dataset, containing the proportion of each grid cell that is covered by protected areas (regardless of its category of protection and not double-counting overlapping conservation status). We then classified each grid cell as belonging to one of the following five classes: 0 (no protection), 0-25%, 25-50%, 50-75%,>75% protected.

Finally, we used the biomes of the world (Figure S1) as made available by (Eric Dinerstein et al., 2017). From these data, we classified each of our 0.25 × 0.25° grid cell as belonging to only one biome, according to the majority class that covered that grid cell. This step allowed us to segment our global analysis and further understand the distribution and trends associated with each biome. All subsequent analyses were performed using the three datasets described above: land use change, protected areas and biomes.

### Land use change analyses

We started our analyses by investigating the coverage of primary and modified areas in the present day (i.e., 2015) at the global scale, per biome and per class of protection. Next, we determined the proportion of primary and modified land that is under protection, as well as the average protection level of the grid cells within each biome. A correlation between the proportion of primary vegetation and proportion of protection was then tested for the hypothesis that higher protection classes would contain higher levels of primary vegetation. Such a hypothesis was assessed both globally and across biomes.

For each one of the SSPs investigated in this study, we assessed how much loss of primary vegetation is projected to occur, globally, per biome and per grid cell from 2015 through 2090, using a decadal interval. Such analysis was performed considering the whole dataset (i.e., regardless of the level of protection), as well as stratified by the five protection classes described before, i.e., to assess whether the loss in primary vegetation across SSPs was significantly different across classes of protection. The significance across biomes and protection classes was assessed using a non-parametric Kruskal-Wallis test and subsequent pairwise comparison Mann–Whitney U-tests, using the Bonferroni correction, where relevant, using the statistical programme R (R Core Team, 2018).

To assess trends over time (from 2015 through 2090 on decadal intervals), we then computed a temporal vector for each grid cell depicting the loss of primary land over time, and implemented a linear regression, accounting for temporal autocorrelation, i.e., using a GLS algorithm, to identify the speed of change associated to each grid cell. Finally, the median slope values of the regressions across SSPs were computed and compared with the values of protection by overlaying the two datasets. A similar procedure was followed to compare the speed of change with original primary vegetation extent at the grid cell level. Moreover, we accumulated the values of change (2015-2090) at the biome, scenario and global scales, to make the same assessment considering the accumulated values, rather than the local (grid cell) values.

## Results

### Distribution of protected areas and primary vegetation areas globally and across biomes

We found that at the global scale by 2015, 14% of the land surface (excluding water bodies) was under some level of protection (Figure 1b, Table 2). Considering cells under protection, on average each grid cell included 16% of protected land (standard error [s.e.] = 0.06%; Figure S2), with a highly skewed distribution of 61% of cells unprotected, 19% with under 25% of the land protected, and only 11% of the grid cells were highly protected (>75%). These proportions varied significantly across biomes (Kruskal-Wallis [KW] test; H = 13,345, p-value <0.001), with the highest protection coverage in Montane Grasslands & Shrublands (27%), Flooded Grasslands & Savannas (25%), and Mangroves (24%) (Table 2). Only six out of the fourteen biomes had a protection coverage above the 17% Aichi Target, with Temperate Grasslands, Savannas & Shrublands being the least protected with only 4% (Table 2). If we analyse the protection of primary vegetation at the grid cell level, we found that the distribution of cells under different levels of protection was highly skewed towards unprotected or low protection (0-25%) globally, with again significant differences across biomes (KW test; H = 13,393, p-value < 0.001, Table 2). In this regard, the maximum proportion of unprotected cells occurred in Deserts & Xeric Shrublands (78%) and the minimum in the Mangroves biome (35%). Contrarily, the highest proportion of highly protected cells (>75%) occurred in the biome Tundra (25%), and the minimum in Temperate Grasslands, Savannas & Shrublands (1%). On average, the highest protection coverage per grid cell was found in the Montane Grasslands & Shrublands (28% ± 0.32, s.e.), and the lowest values were found for Temperate Grasslands, Savannas & Shrublands (4% ± 0.03, s.e.) (Figure S2).

**Table 2.**
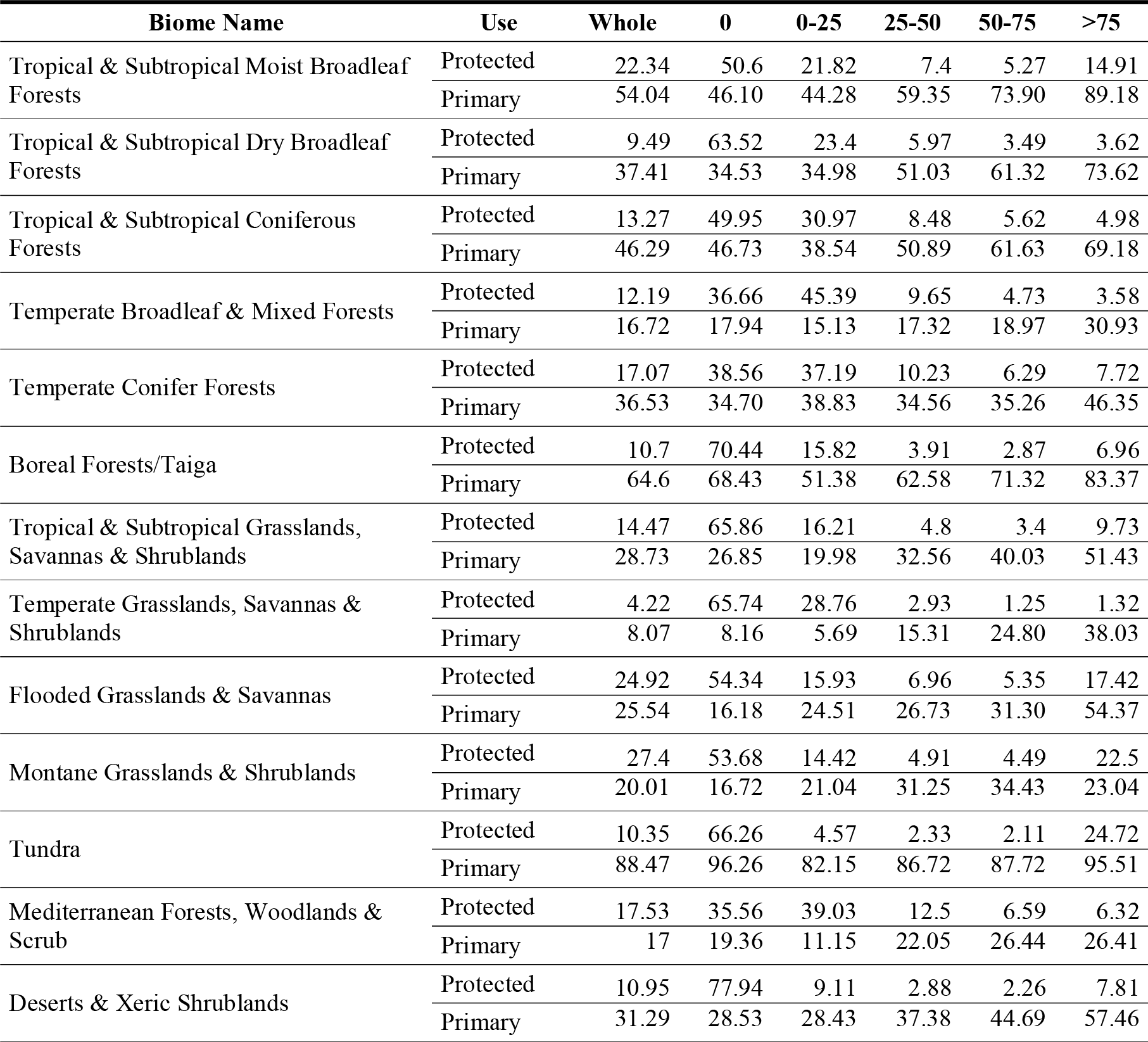
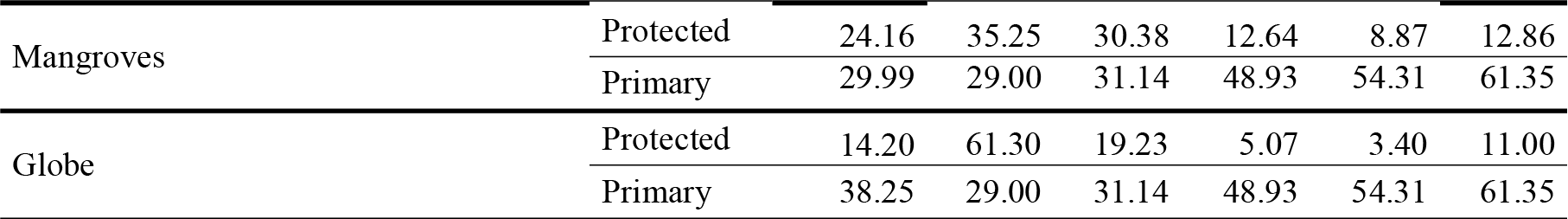
Percentage (%) of biome currently protected or considered primary vegetation, as a whole, as well as considering only the area under different protection classes (from unprotected [0] to more than 75% protected [>75]).

| Biome Name | Use | Whole | 0 | 0-25 | 25-50 | 50-75 | $>75$ |
| --- | --- | --- | --- | --- | --- | --- | --- |
| Tropical & Subtropical Moist Broadleaf Forests | Protected | 22.34 | 50.6 | 21.82 | 7.4 | 5.27 | 14.91 |
|  | Primary | 54.04 | 46.10 | 44.28 | 59.35 | 73.90 | 89.18 |
| Tropical & Subtropical Dry Broadleaf Forests | Protected | 9.49 | 63.52 | 23.4 | 5.97 | 3.49 | 3.62 |
|  | Primary | 37.41 | 34.53 | 34.98 | 51.03 | 61.32 | 73.62 |
| Tropical & Subtropical Coniferous Forests | Protected | 13.27 | 49.95 | 30.97 | 8.48 | 5.62 | 4.98 |
|  | Primary | 46.29 | 46.73 | 38.54 | 50.89 | 61.63 | 69.18 |
| Temperate Broadleaf & Mixed Forests | Protected | 12.19 | 36.66 | 45.39 | 9.65 | 4.73 | 3.58 |
|  | Primary | 16.72 | 17.94 | 15.13 | 17.32 | 18.97 | 30.93 |
| Temperate Conifer Forests | Protected | 17.07 | 38.56 | 37.19 | 10.23 | 6.29 | 7.72 |
|  | Primary | 36.53 | 34.70 | 38.83 | 34.56 | 35.26 | 46.35 |
| Boreal Forests/Taiga | Protected | 10.7 | 70.44 | 15.82 | 3.91 | 2.87 | 6.96 |
|  | Primary | 64.6 | 68.43 | 51.38 | 62.58 | 71.32 | 83.37 |
| Tropical & Subtropical Grasslands, Savannas & Shrublands | Protected | 14.47 | 65.86 | 16.21 | 4.8 | 3.4 | 9.73 |
|  | Primary | 28.73 | 26.85 | 19.98 | 32.56 | 40.03 | 51.43 |
| Temperate Grasslands, Savannas & Shrublands | Protected | 4.22 | 65.74 | 28.76 | 2.93 | 1.25 | 1.32 |
|  | Primary | 8.07 | 8.16 | 5.69 | 15.31 | 24.80 | 38.03 |
| Flooded Grasslands & Savannas | Protected | 24.92 | 54.34 | 15.93 | 6.96 | 5.35 | 17.42 |
|  | Primary | 25.54 | 16.18 | 24.51 | 26.73 | 31.30 | 54.37 |
| Montane Grasslands & Shrublands | Protected | 27.4 | 53.68 | 14.42 | 4.91 | 4.49 | 22.5 |
|  | Primary | 20.01 | 16.72 | 21.04 | 31.25 | 34.43 | 23.04 |
| Tundra | Protected | 10.35 | 66.26 | 4.57 | 2.33 | 2.11 | 24.72 |
|  | Primary | 88.47 | 96.26 | 82.15 | 86.72 | 87.72 | 95.51 |
| Mediterranean Forests, Woodlands & Scrub | Protected | 17.53 | 35.56 | 39.03 | 12.5 | 6.59 | 6.32 |
|  | Primary | 17 | 19.36 | 11.15 | 22.05 | 26.44 | 26.41 |
| Deserts & Xeric Shrublands | Protected | 10.95 | 77.94 | 9.11 | 2.88 | 2.26 | 7.81 |
|  | Primary | 31.29 | 28.53 | 28.43 | 37.38 | 44.69 | 57.46 |
| Mangroves | Protected | 24.16 | 35.25 | 30.38 | 12.64 | 8.87 | 12.86 |
|  | Primary | 29.99 | 29.00 | 31.14 | 48.93 | 54.31 | 61.35 |
| Globe | Protected | 14.20 | 61.30 | 19.23 | 5.07 | 3.40 | 11.00 |
|  | Primary | 38.25 | 29.00 | 31.14 | 48.93 | 54.31 | 61.35 |

**Figure 1.**
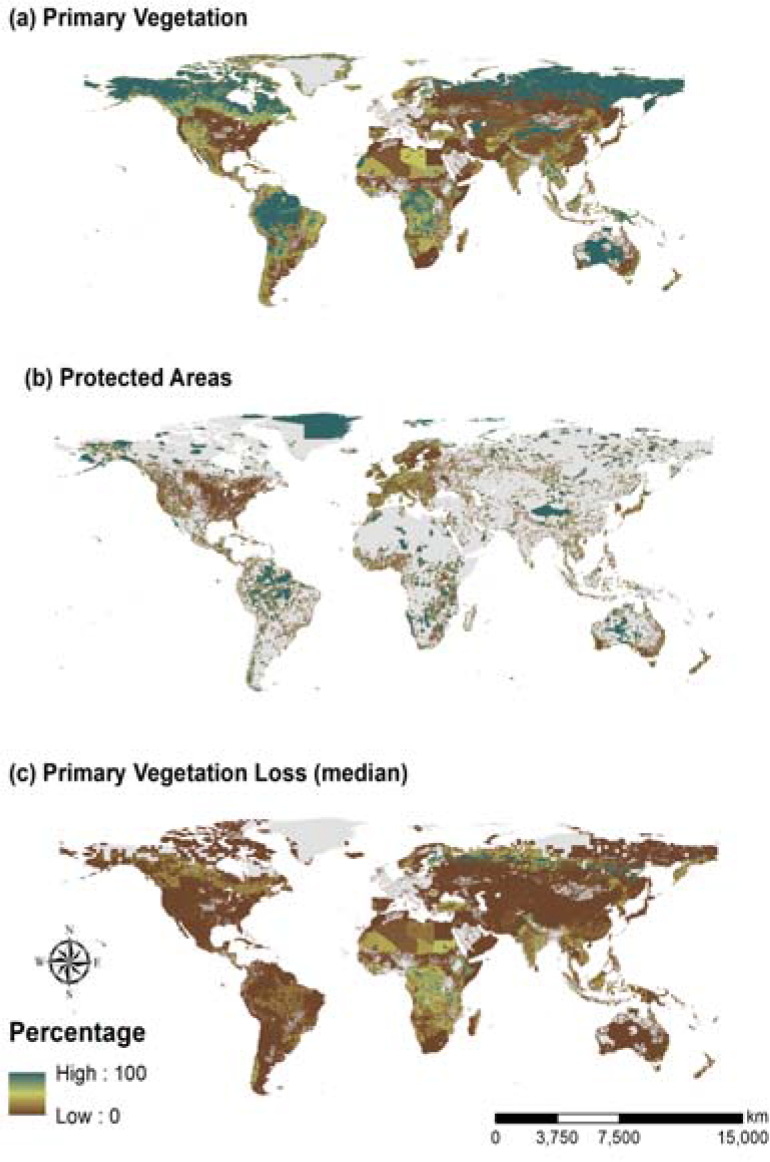
Percentage of the grid cell covered in (a) primary vegetation in 2015, (b) protected area and (c) median loss of primary lands across all SSPs by 2090, relative to 2015 (individual losses per SSP are shown in Figure S3).

Considering our 2015 baseline (Figure 1a), we found that, at the global scale, there was a remaining 38% of areas considered as primary vegetation (forested or non-forested), and 62% of the land had been modified from its natural state. Further, we found a weak positive relationship (t-value = 2.99, p-value = 0.06) between protection level and proportion of natural areas (Table S2), i.e. more natural areas in higher protection cells. At the biome level, there was once again sharp differences, where Temperate Grasslands, Savannas & Shrublands was the biome with the lowest percentage of primary vegetation areas (8%), as opposed to Tundra that was the highest (88%) (Table 2). Within 57% of the biomes, there was indeed a significant linear increase in the proportion of natural areas when considering the protection level (Table S2). However, such coverage varied greatly when analyzed by class of protection (Table 2), both globally and per biome. The average proportion of natural areas per biome varied significantly both without considering the protection level (KW test; H = 35,245, p-value < 0.001), and when considering the cell protection (KW test; H = 57,812, p-value <0.001). In nine out the fourteen biomes, primary vegetation areas were found in greater proportion than modified areas in the highly protected grid cells. On the other hand, in two biomes (Tundra and Boreal Forests/Taiga) primary vegetation areas were observed in higher proportion in unprotected cells.

### Projected changes in primary vegetation areas (2015-2090) globally and per biome

Each of the five scenarios of land use change (SSPs) led to an overall loss of primary vegetation areas from 2015 through 2090 (Figure 2, Table 3). At the global scale, this loss varied between −17.4% in SSP1 to −34.1% in SSP4 (RCP3.4), with an average of −26.84% (2.39% s.e.) across all scenarios (Figure 1c shows median value across all SSPs, whereas Figure S3 shows accumulated change in each individual SSP). Over time, when accumulated globally, the speed of primary vegetation loss (slope of regression, β) is sharper in SSP4 (RCP3.4) and slower in SSP1 (β = −0.45 and β = −0.22, respectively), and the same was observed when considered the local (grid cell average) values (β = −0.50 and β = −0.32, respectively, Figure S4). Further, this loss was higher in pixels with an initial higher proportion of primary vegetation in 2015 (t = 180.03, df = 258,540; p-value < 0.001).

**Figure 2.**
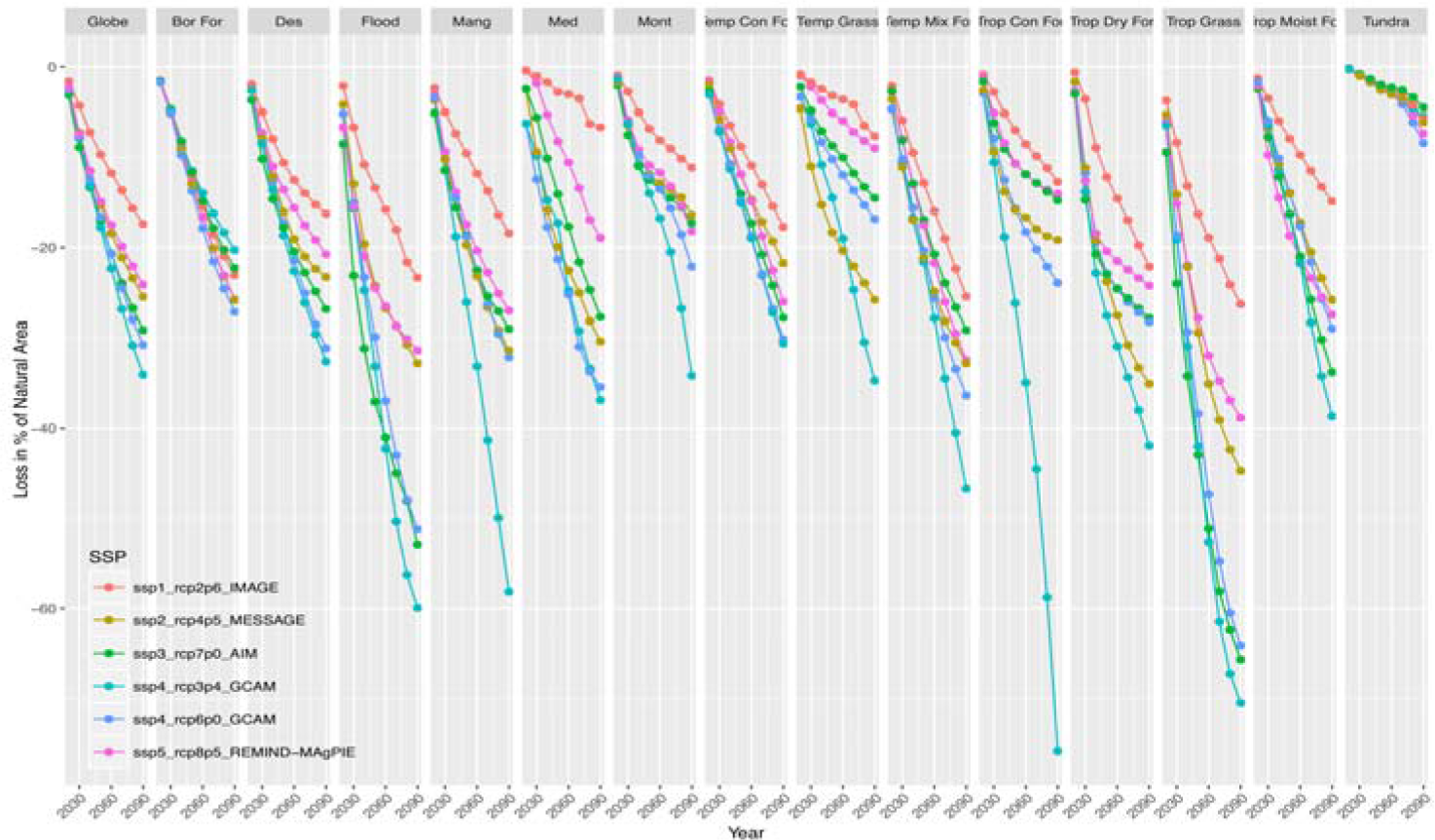
Decadal loss in primary vegetation until 2090, relative to 2015 (in %), globally and per biome, for each of the five land use scenarios (SSPs). Full biome names, Trop Moist For: Tropical & Subtropical Moist Broadleaf Forests; Trop Dry For: Tropical & Subtropical Dry Broadleaf Forests; Trop Con For: Tropical & Subtropical Coniferous Forests; Temp Mix For: Temperate Broadleaf & Mixed Forests; Temp Con For: Temperate Conifer Forests; Bor For: Boreal Forests/Taiga; Trop Grass: Tropical & Subtropical Grasslands, Savannas & Shrublands; Temp Grass: Temperate Grasslands, Savannas & Shrublands; Flood: Flooded Grasslands & Savannas; Mont: Montane Grasslands & Shrublands; Med: Mediterranean Forests, Woodlands & Scrub; Des: Deserts & Xeric Shrublands; Mang: Mangroves.

**Table 3.** Loss in primary vegetation area in each of the land use scenarios, relative to 2015 (in %), per biome and globally.

| Biomes | SSP1 | SSP2 | SSP3 | SSP4a | SSP4b | SSP5 |
| --- | --- | --- | --- | --- | --- | --- |
| Tropical & Subtropical Moist Broadleaf Forests | -14.89 | -25.82 | -33.82 | -38.72 | -28.98 | -27.45 |
| Tropical & Subtropical Dry Broadleaf Forests | -22.13 | -35.11 | -27.82 | -41.93 | -28.27 | -24.25 |
| Tropical & Subtropical Coniferous Forests | -12.74 | -19.16 | -14.77 | -75.74 | -23.94 | -14.03 |
| Temperate Broadleaf & Mixed Forests | -25.44 | -32.82 | -29.14 | -46.72 | -36.41 | -32.47 |
| Temperate Conifer Forests | -17.72 | -21.74 | -27.79 | -30.64 | -30.16 | -26.01 |
| Boreal Forests/Taiga | -23.07 | -25.83 | -22.29 | -20.28 | -27.15 | -25.76 |
| Tropical & Subtropical Grasslands, Savannas & Shrublands | -26.27 | -44.74 | -65.61 | -70.40 | -64.07 | -38.89 |
| Temperate Grasslands, Savannas & Shrublands | -7.65 | -25.81 | -14.51 | -34.75 | -16.84 | -8.99 |
| Flooded Grasslands & Savannas | -23.36 | -32.82 | -52.88 | -59.93 | -51.24 | -31.41 |
| Montane Grasslands & Shrublands | -11.15 | -16.41 | -17.30 | -34.20 | -22.12 | -18.21 |
| Tundra | -5.67 | -6.11 | -4.48 | -5.46 | -8.42 | -7.37 |
| Mediterranean Forests, Woodlands & Scrub | -6.66 | -30.40 | -27.71 | -36.92 | -35.47 | -18.92 |
| Deserts & Xeric Shrublands | -16.25 | -23.28 | -26.83 | -32.63 | -31.15 | -20.75 |
| Mangroves | -18.41 | -31.39 | -29.01 | -58.14 | -32.16 | -27.00 |
| Globe | -17.40 | -25.47 | -29.15 | -34.09 | -30.79 | -24.14 |

We found strong variations across biomes within each scenario (KW test; average H = 54,510, 3596 s.e., p-value < 0.001) and across scenarios within each biome (KW test; average H = 6,664, 2805 s.e., p-value < 0.001). The projected change in primary vegetation across SSPs, varied from a minimum of −76% in SSP4 (RCP3.4) in Tropical & Subtropical Coniferous Forests, Savannas & Shrublands to a maximum of −4.5% in SSP3 in Tundra (Table 3). On average, Tundra is the least impacted biome (−6.25%, 0.58 s.e.), whereas Tropical & Subtropical Grasslands, Savannas & Shrublands is the highest impacted biome (−51.7%, 7.2 s.e.). As expected, both globally and in all but two biomes (Tundra and Boreal Forests/Taiga), SSP1 was the least harmful scenario, and interestingly, SSP1 was not the best scenario for the two most highly protected biomes (Tundra and Boreal Forests), where SSP4 (RCP3.4) led to fewer losses (Figure S3 and S4).

### Projected changes in natural areas (2015-2090) globally and per biome considering protection

When considering the protection level of each grid cell we found that the areas under greatest threat of conversion are mostly located in the unprotected and 0-25% categories (Figure 3), although there was still a large proportion of change in the highly protected areas (varying from −18% to −30%, in SSP1 and SSP4a, respectively). Further, there was no significant correlation found between protection coverage and average proportion of vegetation loss (t = 1.83, df = 258,540; p-value = 0.07).

**Figure 3.**
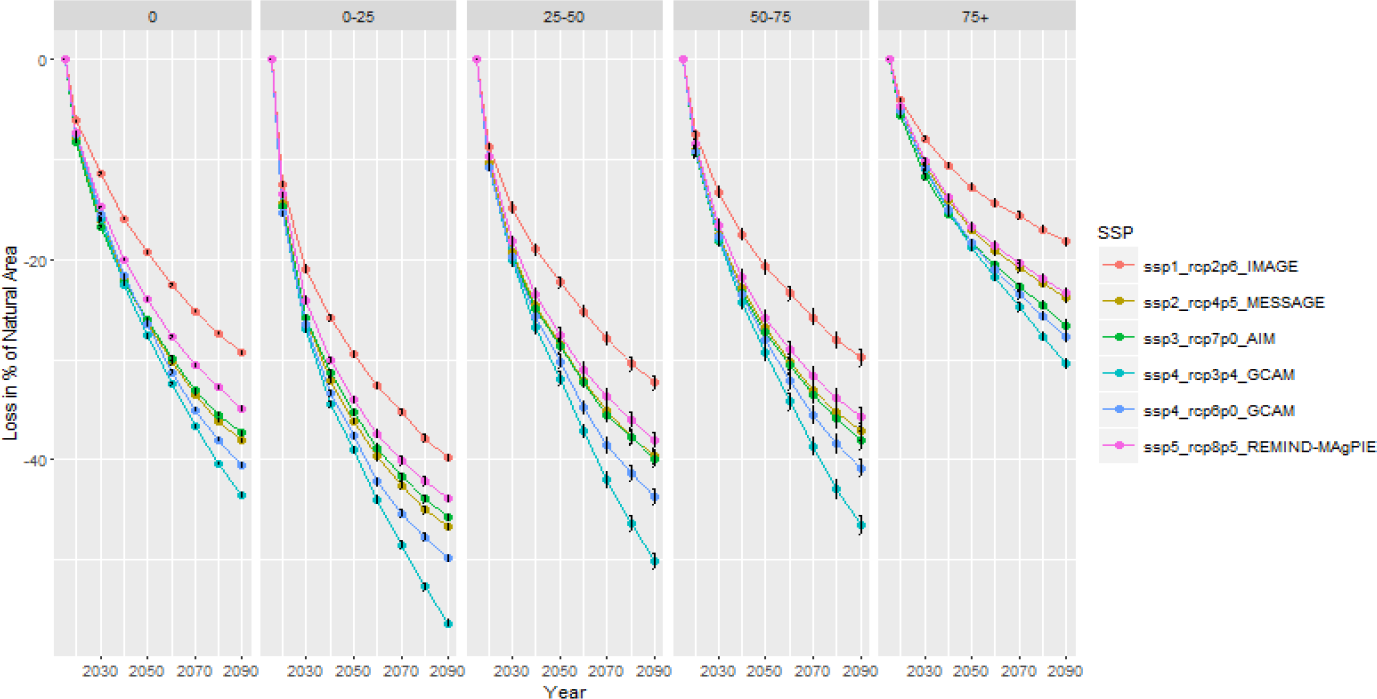
Decadal average loss until 2090 (relative to 2015 in %) within each scenario of land use change (SSPs) considering the protection coverage of each grid cell.

When averaging the overall change between 2015 and 2090 (across all scenarios), we found significant differences across biomes and protection level (Table S3). In detail, in the majority of the biomes the protection class with the highest projected loss in primary vegetation is either unprotected (in 7 out of 14 biomes) or low protection (0-25%, in 5 out of 14 biomes). In the Mediterranean Forests, Woodlands & Scrub the largest projected loss occurred in the highest protected grid cells, despite comprising the lowest proportion of cells in the Biome with only 6.32% of the grid cells falling in this protection category (Table 2).

Finally, in order to highlight areas for intervention to prevent projected losses from occurring, we overlapped the overall (and trend) in projected primary vegetation loss (2015-2090), with the protection class (Figure 4). We found that the Tropical Forests in Central Africa and the Boreal Forests of North Euro-Asia and Canada emerge as the areas where most projected change occurs in areas where existing protection coverage is still low. Similarly, areas in India and Southeast Asia emerge as potential areas for intervention as they have significant projected loss of primary vegetation, and considerably low (0-25%) protection.

**Figure 4.**
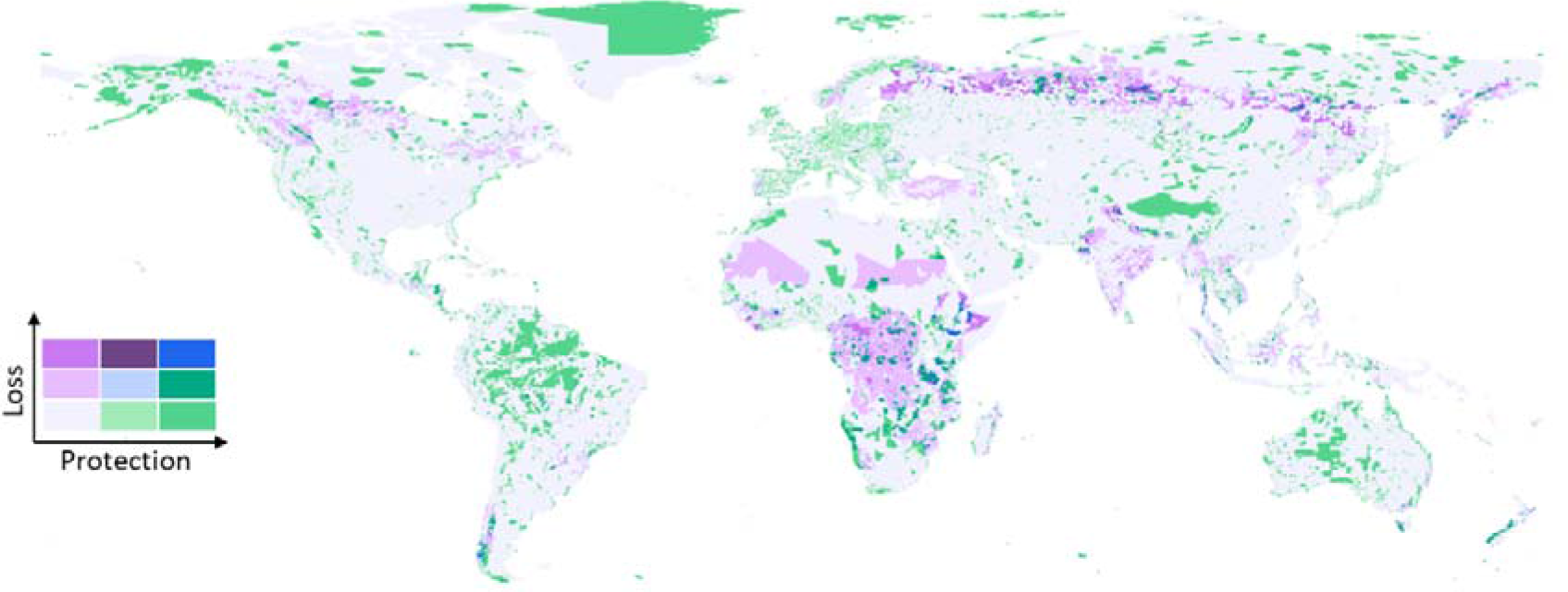
Projected primary vegetation loss (median across SSPs, individual results for each SSP in Figure S5) from 2015 through 2090, overlapped with proportion of protected (0-25%, 25-75%, >75%).

## Discussion

Despite international conservation efforts, particularly in relation to the expansion of protected areas worldwide (Thomas & Gillingham, 2015), we have been unable to slow down the destruction of natural habitats, as recently highlighted by the IPBES Global Assessment (Díaz et al., 2019) and the near real time monitoring platform for forests, Global Forest Watch (Curtis, Slay, Harris, Tyukavina, & Hansen, 2018). One of the key elements of biodiversity targets is the ability to preserve environmental representativeness, which has not driven protected area expansion, with the focus placed on factors such as low productive value, population and tourism (Baldi et al., 2017). The presence of natural areas (primary vegetation) was highly skewed towards certain biomes (most under the desired 17% protected coverage Aichi Target), and according to the modelled data used in our study, the weak relationship between the extent of the remaining natural areas and the extent of protection across biomes, suggests that we are endangering the representativeness of all biomes (as desired by the Aichi targets). These regions include some of the most biologically distinctive, species-rich ecosystems on Earth, such as tropical forests, thus compromising the preservation of genetic resources from a wide variety of life on Earth. Further, as highly protected cells tended to contain larger proportions of natural areas, the remaining natural areas of the world are becoming confined to current protected areas. This pattern highlights the need to ensure the efficacy of these areas in preventing further degradation, which has not always been the case (e.g., Rosa, Rentsch, & Hopcraft, 2018; Soares-Filho et al., 2010). Further, there is a dichotomy between proportion of area covered and ‘connectivity’ of the protected areas network, for instance, biomes such as Temperate Grasslands, Savannas & Shrublands emerged as having a high proportion of coverage (almost a quarter), but very fragmented, with a low proportion of full protected grid cells (1%), suggesting low connectivity (Saura et al., 2017).

As we essentially failed to achieve the targets proposed by the CBD by 2020 (Amengual & Alvarez-Berastegui, 2018), the new conservation agenda, at the global scale, is under discussion, with a great focus on restoring degraded ecosystems. For instance, the UN declared 2021-2030 as the Decade for Ecosystem Restoration, and recent studies (Bastin et al., 2019) state that planting forests (afforestation) would be the cheapest solution to address climate change. Nevertheless, it is critical to aid restoration with the preservation of the remains of natural vegetation as these contain the highest biodiversity levels (Newbold et al., 2015), genetic diversity, bank seeds, even in small patches (Wintle et al., 2019). Independently of the scenario followed, the current human development trajectories all lead to further primary vegetation loss. Despite numerous studies drawing attention to the disparities in habitat loss and protection (Hoekstra et al., 2005), and showing that halting agricultural expansion, increasing agriculture efficiency, shifting diets and reducing waste (Foley et al., 2011; Lambin & Meyfroidt, 2011), would greatly help preserve existing habitats, the climate change community still largely ignores these aspects in their ‘most positive’ views of the world. Moreover, the recent IPBES call for transformative change in our society to preserve global biodiversity, make these novel visions (Rosa et al., 2017) influencing human development critically needed for our sustainability. In this context, our results show that even under the best possible scenario (SSP1) we will continue the ‘anthropogenization’ of our world (Ellis et al., 2010). This means that further biodiversity loss is unavoidable unless we act now to prevent further expansion of land use into natural ecosystems (Pouzols et al., 2014).

Serious efforts to conserve the remaining 38% of natural areas need to target regions of the world where land use change is expected to happen, thus avoiding or minimizes the chances of that change to occur (pro-active rather than reactive conservation). On the one hand, tropical forests in Central Africa and Southeast Asia, as well as natural vegetation in India, emerge as highly likely to be destroyed (under all scenarios) and where protection coverage is still low. As land use is a highly locked-in process (Guerra et al. under review), i.e. once it changes it rarely reverses, this is the moment to rally internationally, support these nations, and act before we lose these amazingly rich biodiversity hotspots. On the other hand, Boreal forests, which still have low protected coverage (11%), are likely to undergo extensive land use change particularly under more ‘aggressive’ scenarios. Such areas may experience even more important biological loss under the context of climate change, with impact on species distribution (Tuanmu et al., 2013) and on carbon sequestration (Melillo et al., 2016).

Recent calls for more ambitious conservation targets (Mace et al. 2018), including to protect half of the Earth’s land area (Dinerstein et al., 2019; Dinerstein et al., 2017), seem unlikely under the projected changes and given that we failed to achieve existing ones. This is further highlighted by our inaction to address head-on the issue of feeding a growing population with current dietary requirements (Mehrabi et al. 2018) or the teleconnections of dispersed impacts between regions of the globe (Marques et al., 2019). More than defining new area-based targets, a new paradigm that explicitly connects targets with indicators of desired conservation outcomes (Barnes et al., 2018) needs to account for the expected conflict between land use change (Wolff et al. 2018), protection of remaining native vegetation, and restoration of degraded ecosystems under climate change. Apart from improving the efficacy of existing protected areas, new conservation and restoration mechanisms need to be developed to address this wicked challenge. Independently, proactive conservation of the remaining natural vegetation is key to ensure the preservation of biological diversity, aid the recovery of degraded habitats, and help to mitigate climate change.

