## Supporting Information for "Pathways of human development threaten biomes’ protection and their remaining natural vegetation"

**Tables**

**Table S1 –** Original land use classes of the Land Use Harmonized dataset (http://luh.umd.edu/) and classification used in this study.

| **Original land use class name** | **New class** |
| --- | --- |
| Forested primary land (primf) | Primary |
| Non-forested primary land (primn) | Primary |
| Potentially forested secondary land (secdf) | Modified |
| Potentially non-forested secondary land (secdn) | Modified |
| Managed pasture (pastr) | Modified |
| Rangeland (range) | Modified |
| Urban land (urban) | Modified |
| C3 annual crops (c3ann) | Modified |
| C3 perennial crops (c3per) | Modified |
| C4 annual crops (c4ann) | Modified |
| C4 perennial crops (c4per) | Modified |
| C3 nitrogen-fixing crops (c3nfx) | Modified |

**Table S2 –** Linear regression between proportion of primary vegetation and protection of protection per biome and globally.

| **Biome Name** | **Estimate** | **Std. Error** | **t value** | **p-value** | **R^2^** |
| --- | --- | --- | --- | --- | --- |
| Tropical & Subtropical Moist Broadleaf Forests | 11.555 | 1.939 | 5.958 | 0.009 | 0.922 |
| Tropical & Subtropical Dry Broadleaf Forests | 10.406 | 1.365 | 7.622 | 0.005 | 0.951 |
| Tropical & Subtropical Coniferous Forests | 6.788 | 2.061 | 3.293 | 0.046 | 0.783 |
| Temperate Broadleaf & Mixed Forests | 3.257 | 1.550 | 2.101 | 0.126 | 0.595 |
| Temperate Conifer Forests | 2.051 | 1.444 | 1.420 | 0.251 | 0.402 |
| Boreal Forests/Taiga | 4.906 | 3.179 | 1.543 | 0.220 | 0.443 |
| Tropical & Subtropical Grasslands, Savannas & Shrublands | 6.938 | 1.925 | 3.603 | 0.037 | 0.812 |
| Temperate Grasslands, Savannas & Shrublands | 7.581 | 1.564 | 4.846 | 0.017 | 0.887 |
| Flooded Grasslands & Savannas | 8.219 | 2.116 | 3.885 | 0.030 | 0.834 |
| Montane Grasslands & Shrublands | 2.579 | 2.198 | 1.174 | 0.325 | 0.315 |
| Tundra | 0.383 | 2.248 | 0.170 | 0.876 | 0.010 |
| Mediterranean Forests, Woodlands & Scrub | 2.944 | 1.560 | 1.887 | 0.156 | 0.543 |
| Deserts & Xeric Shrublands | 7.405 | 1.266 | 5.849 | 0.010 | 0.919 |
| Mangroves | 8.483 | 1.764 | 4.809 | 0.017 | 0.885 |
| Globe | 7.857 | 2.624 | 2.995 | 0.06 | 0.67 |

**Table S3 –** Loss of primary vegetation by 2090 (in % relative to 2015) per grid cell in all five scenarios, considering the protection coverage of each grid cell.

| **Biome** | **0** | | **0-25** | | **25-50** | | **50-75** | | **>75** | |
| --- | --- | --- | --- | --- | --- | --- | --- | --- | --- | --- |
|  | **Mean** | **Std** | **Mean** | **Std** | **Mean** | **Std** | **Mean** | **Std** | **Mean** | **Std** |
| Boreal  Forests/Taiga | -38.59 | 2.62 | -53.86 | 1.99 | -42.89 | 2.62 | -35.52 | 2.24 | -27.45 | 2.53 |
| Deserts &  Xeric Shrublands | -42.14 | 6.55 | -29.06 | 5.93 | -26.36 | 6.17 | -28.18 | 5.85 | -31.67 | 5.36 |
| Flooded Grasslands  & Savannas | -61.67 | 7.60 | -49.81 | 9.32 | -55.68 | 8.02 | -58.86 | 7.92 | -60.30 | 17.49 |
| Mangroves | -55.22 | 16.01 | -46.20 | 11.22 | -31.85 | 11.02 | -36.32 | 11.12 | -35.28 | 9.36 |
| Mediterranean Forests,  Woodlands & Scrub | -59.75 | 12.21 | -41.33 | 11.37 | -36.82 | 10.82 | -46.37 | 9.24 | -63.61 | 5.39 |
| Montane Grasslands  & Shrublands | -23.56 | 8.86 | -41.54 | 7.05 | -28.70 | 7.47 | -27.91 | 8.35 | -12.95 | 4.56 |
| Temperate Broadleaf  & Mixed Forests | -42.18 | 9.37 | -60.48 | 5.72 | -57.90 | 6.72 | -65.82 | 5.15 | -61.80 | 5.10 |
| Temperate Conifer Forests | -49.83 | 6.46 | -43.11 | 3.25 | -47.36 | 3.84 | -46.92 | 4.27 | -40.00 | 3.60 |
| Temperate Grasslands,  Savannas & Shrublands | -21.80 | 9.79 | -38.86 | 7.98 | -28.24 | 8.42 | -25.12 | 10.97 | -21.16 | 10.18 |
| Tropical & Subtropical  Coniferous Forests | -32.14 | 22.10 | -42.80 | 18.58 | -33.56 | 20.78 | -28.72 | 22.67 | -22.04 | 23.12 |
| Tropical & Subtropical  Dry Broadleaf Forests | -46.46 | 8.14 | -40.71 | 10.82 | -42.64 | 12.22 | -40.09 | 11.14 | -23.94 | 6.58 |
| Tropical & Subtropical  Grasslands, Savannas & Shrublands | -66.33 | 11.23 | -62.59 | 9.33 | -61.34 | 11.94 | -60.60 | 11.75 | -61.39 | 15.12 |
| Tropical & Subtropical  Moist Broadleaf Forests | -52.09 | 9.51 | -49.26 | 7.91 | -42.27 | 7.75 | -34.94 | 6.99 | -18.76 | 4.04 |
| Tundra | -5.91 | 1.30 | -16.03 | 5.19 | -12.38 | 3.00 | -11.55 | 2.49 | -3.94 | 0.65 |

**Figures**

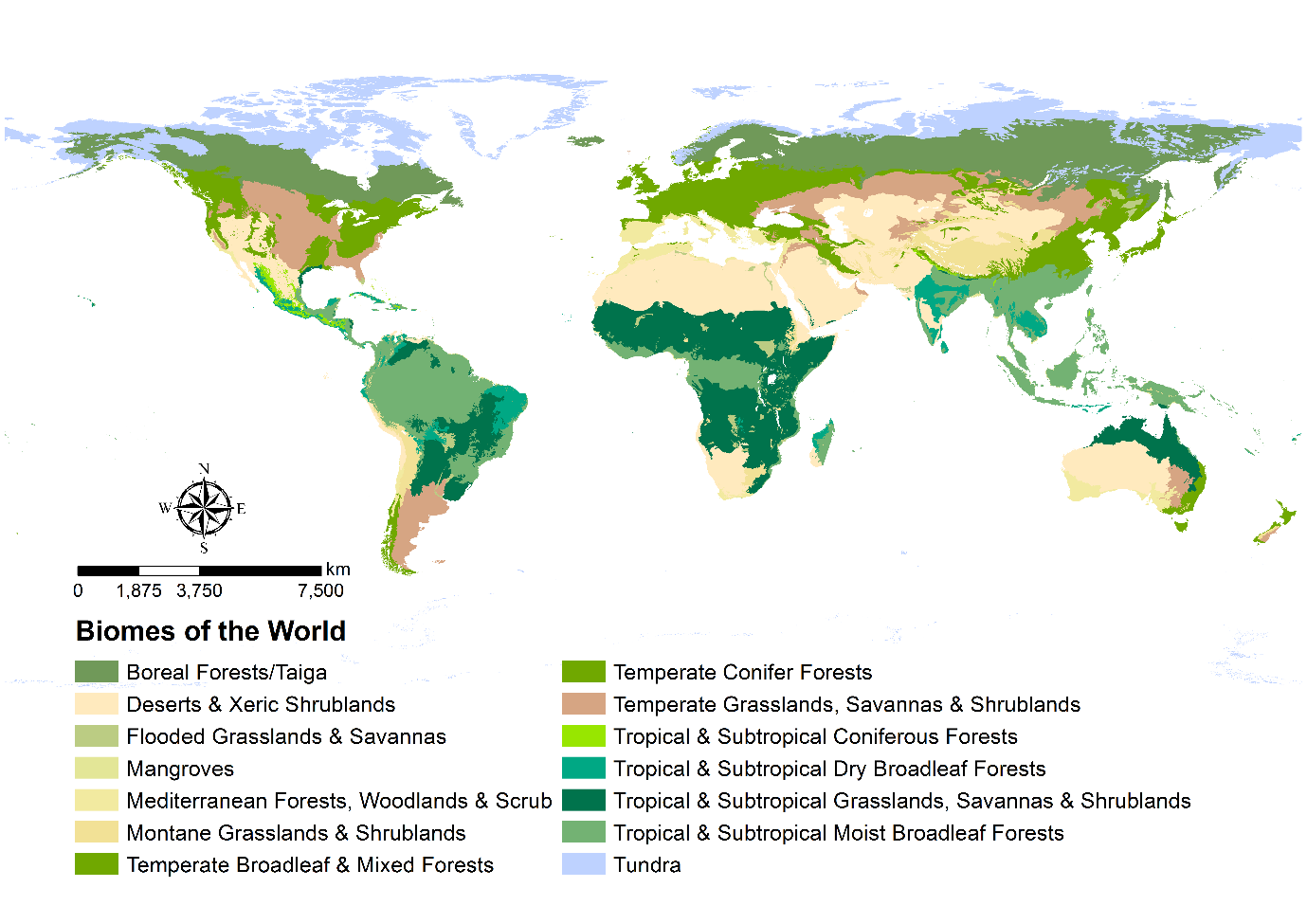

**Figure S1 –** Biomes of the world (original data provided by Dinerstein et al., (2017)).

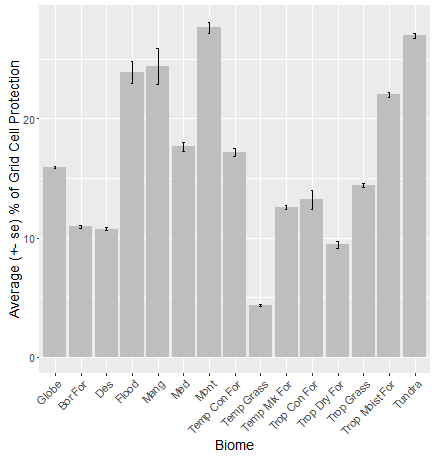

**Figure S2 –** Average protection coverage (in %) within biomes and globally. Full biome names, Trop Moist For: Tropical & Subtropical Moist Broadleaf Forests; Trop Dry For: Tropical & Subtropical Dry Broadleaf Forests; Trop Con For: Tropical & Subtropical Coniferous Forests; Temp Mix For: Temperate Broadleaf & Mixed Forests; Temp Con For: Temperate Conifer Forests; Bor For: Boreal Forests/Taiga; Trop Grass: Tropical & Subtropical Grasslands, Savannas & Shrublands; Temp Grass: Temperate Grasslands, Savannas & Shrublands; Flood: Flooded Grasslands & Savannas; Mont: Montane Grasslands & Shrublands; Med: Mediterranean Forests, Woodlands & Scrub; Des: Deserts & Xeric Shrublands; Mang: Mangroves.

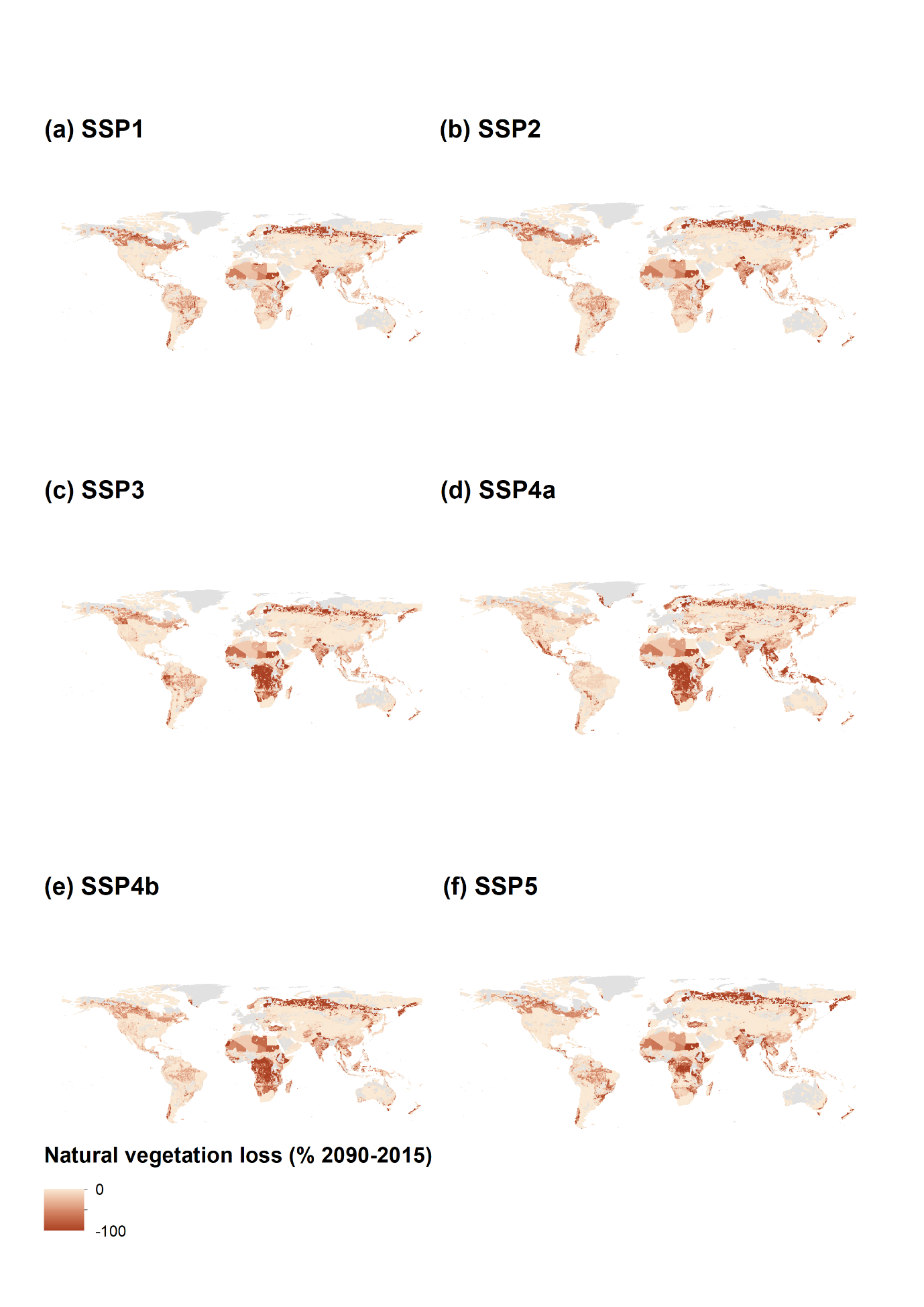

**Figure S3** – Accumulated primary vegetation loss by 2090 (in % relative to 2015), for each individual Shared Socio-Economic Pathway (SSP).

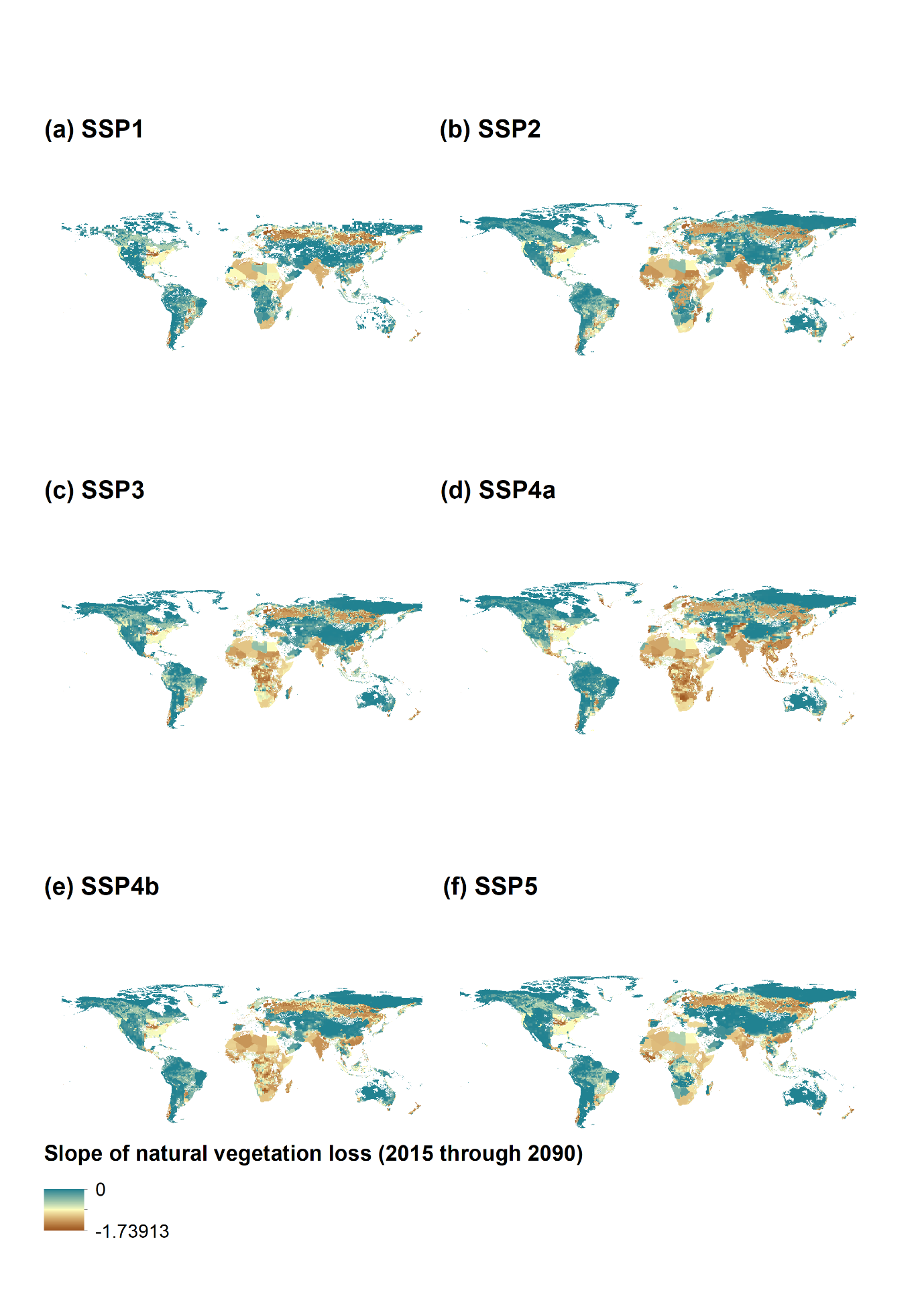

**Figure S4** – Slope of generalized least squares regression for projected primary vegetation loss (from 2015 through 2090), for each individual Shared Socio-Economic Pathway (SSP).
